# SUMO proteases SPF1/2 inhibit seed filling and fatty acid biosynthesis by deSUMOylating WRI1

**DOI:** 10.1101/2025.08.01.668025

**Authors:** Ruihua Huang, Mengrui Wen, Yangyang Chen, Bojing Feng, Miaolun Yuan, Chao Yang, Huizi Huang, Jianhai Mo, Qiuchan Lu, Wenjuan Lai, Zixuan Ma, Lin Zhang, Hongqing Li, Shengchun Zhang

## Abstract

Seed filling and fatty acid biosynthesis are critical processes for seed development and oil accumulation in plants, regulated by intricate post-translational mechanisms. Here, we elucidate the molecular mechanism by which SUMO proteases SPF1 and SPF2 regulate seed filling and fatty acid biosynthesis by modulating the stability of the WRI1 protein through deSUMOylation in *Arabidopsis*. SPF1 and SPF2 were found to negatively regulate seed filling and oil accumulation by promoting WRI1 degradation via deSUMOylation, which enhances its interaction with BPM proteins and subsequent ubiquitination. Our findings establish a dynamic interplay between deSUMOylation and ubiquitination in controlling WRI1 stability, unveiling a novel mechanism for seed filling. These findings establish SPF1/2 as crucial factor in maintaining WRI1 stability and seed filling, providing valuable genetic resources and a theoretical foundation for improving seed quality and oil content in agricultural production.

## Introduction

WRINKLED1 (WRI1) is an APETALA2 (AP2) transcriptional factor that directly or indirectly regulate genes involved in glycolysis and fatty acid (FA) biosynthesis. (Cernac & Benning, 2004; Maeo et al., 2009; Ma et al., 2013). Mutations in WRI1 result in a severe reduction in seed oil content, by approximately 80% (Baud et al., 2007). Conversely, overexpression of WRI1 leads to a significant increase in the expression of genes involved in glycolysis and fatty acid synthesis, as well as an elevation in oil content in seeds (Cernac & Benning, 2004; Sanjaya et al., 2011). The regulation of WRI1 involves post-translational modifications and transcriptional control by molecular chaperones. KIN10 interacts with and phosphorylates WRI1, leading to its proteasomal degradation (Zhai et al., 2017). Trehalose-6-phosphate, an effective inhibitor of KIN10, stabilizes WRI1 protein and promotes lipid biosynthesis in seeds and nutrient tissues (Zhan et al., 2018; Zhai et al., 2021). Additionally, BPMs proteins, the CULLIN3-based E3 ligase adaptor protein, bind to WRI1, promoting its proteasome degradation (Chen et al., 2013). Recently, we reported that the Pumilio RNA binding protein APUM24 seems to lower the abundance of BPM mRNA so as to stabilize the WRI1(Huang et al., 2021). Moreover, SUMO E3 ligase SIZ1 maintains the stability of WRI1 protein through SUMOylation under high temperature stress, promoting seed filling and fatty acid content (Huang et al., 2025). Furthermore, MED15, TCP4, BLISTER and CDK8 have been shown to play vital roles in the regulation of downstream target genes of WRI1 (Kim et al., 2016; Kong et al., 2020; Huang et al., 2022; Zhai et al., 2023).

SUMOylation, involving the attachment of the small ubiquitin-related modifier (SUMO) to targeted proteins via transfer of the SUMO molecular to lysine residues, represents a reversible and essential post-translational modification prevalent in eukaryotic cells, especially under stress conditions (Yoo et al., 2006; Augustine & Vierstra, 2018). This modification can significantly alter protein function, activity, localization, and stability (Wilkinson & Henley, 2010; Jentsch & Psakhye, 2013; Augustine & Vierstra, 2018). Typically, the SUMO molecule is covalently linked to a substrate protein at a lysine residue within a consensus motif via its C-terminal. This process is sequentially catalyzed by an E1 activating enzyme complex, an E2 conjugating enzyme, and usually an E3 ligase (Han et al., 2021). SUMOylation modulates substrate protein subcellular localization, stability, activity, and interactions with other proteins, thereby playing a pivotal role in diverse plant biological processes. These include stress response (Yu et al., 2024; Zheng et al., 2022), immune defense (Liu et al., 2023; Niu et al., 2019; Zhang et al., 2024), hormone signaling (Zhang et al., 2023; Srivastava et al., 2022), flowering regulation (Jin et al., 2008), seed filling (Huang et al., 2025), photomorphogenesis (Lin et al., 2016; Xiong et al., 2023), and nutrient acquisition (Miura et al., 2005; Park et al., 2011).In contrast, SUMOylation is counteracted by SUMO-specific proteases (Castro et al., 2018). The Arabidopsis genome is presumed to encode eight ubiquitin-like proteases (ULPs), including OTS1, OTS1, ESD4, SPFs and ULP1a (Castro et al., 2018). Several target substrates of ULPs that have been identified in the regulation of plant development and stress responses. For instance, ESD4 controls STOP1 SUMOylation, enhancing STOP1 association with the AtALMT1 promoter to regulate Al resistance in Arabidopsis (Fang et al., 2020). OTS1 and OTS2 modulate the protein abundance of SUMO conjugates in salt stress responses, partly through deSUMOylation and destabilization of growth-repressing DELLA proteins (Conti et al., 2008; Conti et al., 2014), and they also control light-induced signaling by deSUMOylating PHYB (Sadanandom et al., 2015). Additionally, OTS1 and OTS2 regulate jasmonic acid signaling by altering the abundance of JAZ protein through deSUOMOylation (Srivastava et al., 2018). Recently, ULP1a has been found to deSUMOylated BZR1, attenuating BR-promoted growth during salt stress in Arabidopsis (Srivastava et al., 2020). Furthermore, SPF1 and SPF2 regulate the stability of FLC and EDA9, thereby affecting flowering time and fertility, partially through deSUOMOylation (Kong et al., 2017; Liu et al., 2017). SPF1 and SPF2 also control WRKY33 SUMOylation in response to plant immunity (Verma et al., 2021).

This study elucidates the molecular mechanism by which the SUMO proteases SPF1 and SPF2 regulate seed filling and fatty acid accumulation by modulating the stability of the WRI1 protein via deSUMOylation. Specifically, SPF1 and SPF2 mediate the deSUMOylation of WRI1, thereby enhancing its interaction with BPMs proteins and promoting WRI1 degradation through ubiquitination. Our findings not only reveal a novel post-translational modification mechanism for WRI1 but also uncover a new regulatory pathway for seed filling, mediated by SPF1/2-dependent deSUMOylation of WRI1.

## Results

### SPF1 and SPF2 affect seed filling and oil accumulation

The SUMO proteases SPF1 and SPF2 affect seed development (Liu et al., 2017). To investigate the role of SPF1 and SPF2 in seed filling, we initially obtained T-DNA insertion mutants *spf1-1 (SALK_040576C)* and *spf2-1 (SALK_023493C)*, subsequently generating the *spf1-1spf2-1* double mutant through genetic crossing. Phenotypic analysis revealed that Consistent with previous studies, the *spf1-1spf2-1* double mutant produced significantly plumper and heavier seeds compared to wild-type (Fig.1A-1C). Meanwhile, the *spf1-1* and *spf2-1* single mutants also exhibited slightly larger and heavier seeds than WT seeds (Fig.1A-1C). Since seed filling mainly involves the storage accumulation process, we further examined the seed mass and oil content of the *spf1*, *spf2*, and *spf1-1spf2-1* mutants. The total fatty acid content and seed mass of *spf1-1* and *spf2-1* seeds were slightly increased, whereas the *spf1-1spf2-1* double mutant showed an approximate 15% and 30% increase respectively compared to WT (Fig.1B, 1C). Additionally, there was no difference in fatty acid composition in *spf1-1* and *spf2-1* mutants(data not shown) These results suggest that SPF1 and SPF2 may influence seed filling. To validate this hypothesis, we analyzed the seed filling progress at various stages from 4 days after pollination (DAP) to 16 DAP. The seed filling process was notably accelerated at 10 DAP in *spf1-1* and *spf2-1* mutants, especially in the *spf1*-*1spf2*-*1* double mutant compared to WT (Fig. 1D), indicating that SPF1 and SPF2 might act as negative regulators of seed filling.

**Figure 1.**
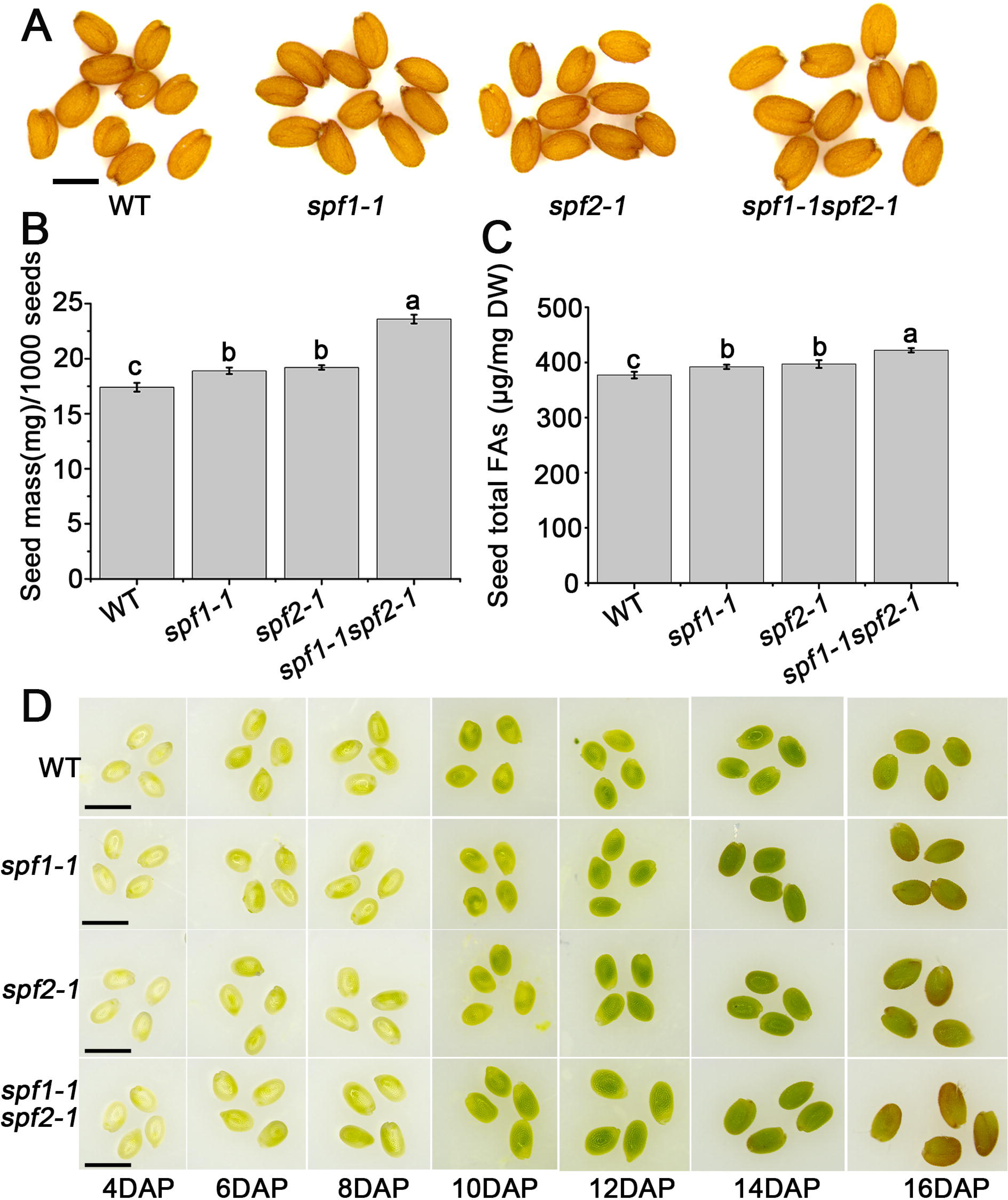
SPF1 and SPF2 affect seed filling and oil accumulation. (A). Morphological comparison of mature seeds from WT, *spf1-1*, *spf2-1*, and *spf1-1spf2-1* plants. Bars = 500 μm. (B). Seed mass comparison among WT, *spf1-1*, *spf2-1*, and *spf1-1spf2-1* plants. Data are presented as means ± standard deviation (SD) (N = 3). Seed mass values labeled with the same letter are not significantly different, as determined by Tukey’s HSD test (P< 0.05). (C) Fatty acid contents in seeds of WT, *spf1-1*, *spf2-1*, and *spf1-1spf2-1* plants. Data are presented as means ± standard deviation (SD) (N = 3). Fatty acid contents sharing the same letter are not significantly different, as determined by Tukey’s HSD test (P< 0.05). (D) Observation of the filling process of *spf1-1*and *spf2-1* seeds, Bars= 1 mm. DAP, days after pollination.

### SPF1 and SPF2 are expressed during seed filing

To determine whether SPF1 and SPF2 participate in regulating seed filling, we examined their expression patterns during seed filling. Reverse transcription quantitative PCR (RT-qPCR) revealed that *SPF1* expression remained relatively low from 2 to 8 days after pollination (DAP) but increased from 10 DAP. This was corroborated by the pSPF1:β-glucuronidase (GUS) reporter assay (Fig. 2A, 2B). Similarly, *SPF2* expression in embryos was lower but exhibited a comparable pattern (Fig. 2C, 2D). These results suggest that both *SPF1* and *SPF2* are expressed during seed filling. To assess whether SPF1 and SPF2 protein levels accumulate during seed filling, we generated *pSPF1:SPF1-GFP* and *pSPF2:SPF2-GFP* plants. GFP signals indicated that SPF1 and SPF2 accumulate in embryonic cells from 6 to 16 DAP, with both proteins localized in the nucleus of embryonic cells (Fig. 2E–2H). Furthermore, to determine whether SPF1 and SPF2 exhibit similar protein expression patterns to WRI1 during filling, we analyzed the accumulation of SPF1, SPF2, and WRI1 proteins in seeds at different filling stages using western blotting. Results showed significant expression of SPF1, SPF2, and WRI1 proteins in seeds at 8, 10, 12, and 14 DAP (Fig. 2I, 2J). Notably, the expression of SPF1 and SPF2 overlapped with that of WRI1 primarily during the 8–14 DAP stages, indicating their potential roles in seed filling.

**Figure 2.**
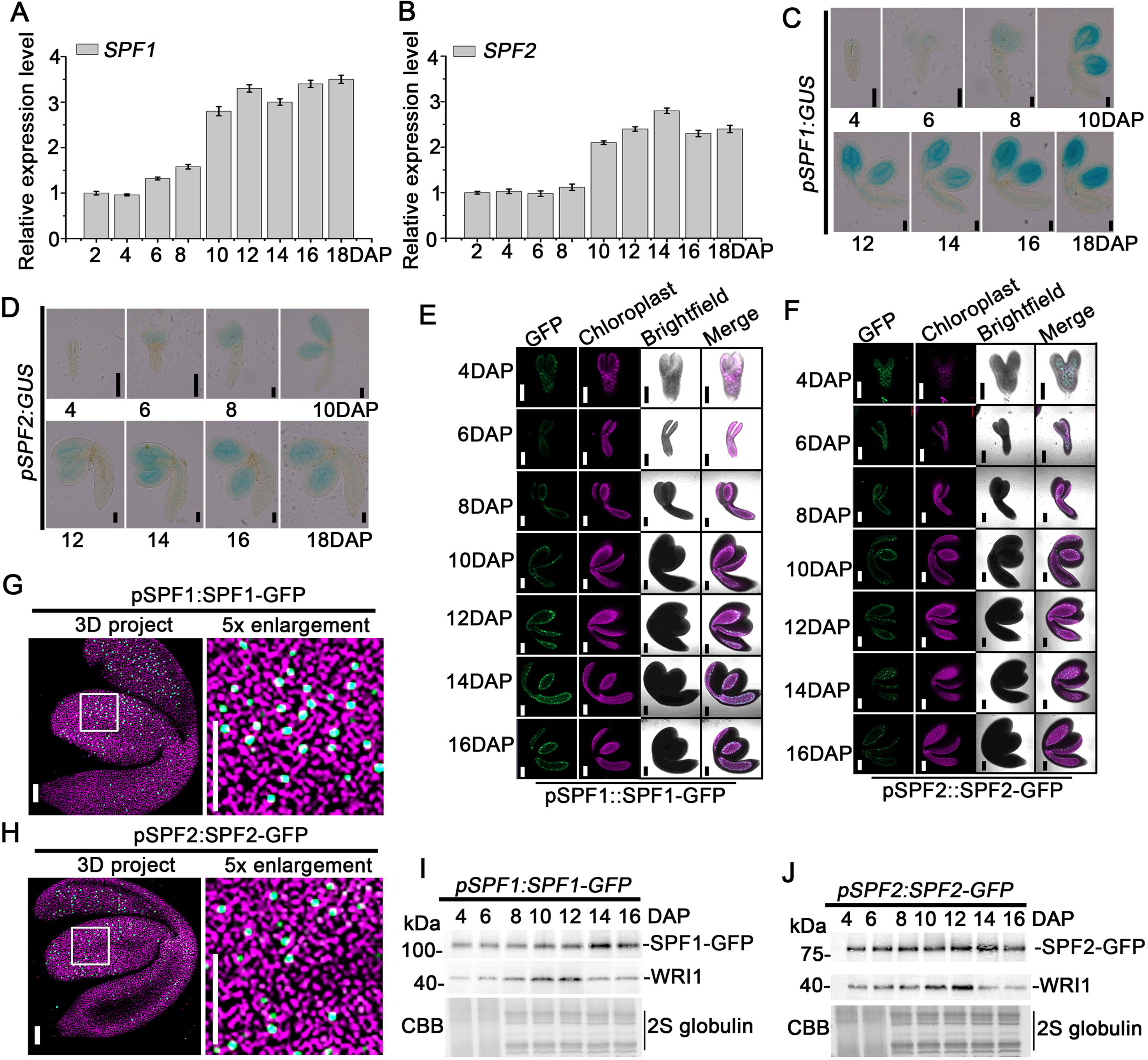
SPF1 and *SPF2* are expressed during seed filling. (A-B) Detection of SPF1 and SPF2 expression levels in *Arabidopsis* developing seeds using RT-qPCR. DAP, days after pollination. Data shown are means ± standard deviation (SD) (N = 3). *UBQ10* was used as a reference gene. (C-D) Visualization of SPF1 and SPF2 expression pattern in *Arabidopsis* developing seeds through GUS staining. DAP, days after pollination. Bars=200 μm. (E-F) Accumulation pattern of SPF1 and SPF2 protein during embryonic development. GFP fluorescence (green) was observed in *pSPF1:SPF1-GFP* or *pSPF2:SPF2-GFP* embryos. Chloroplast, chloroplast fluorescence (magenta), Merge, overlap of SPF1-GFP, SPF2-GFP and chloroplast fluorescence by brightfield microscopy. Bars= 100 μm. (G-H) Enlarged view of SPF1 and SPF2 protein localization in embryonic cells at 12 DAP under normal temperature. Bars= 50 μm. (I) Protein levels of SPF1 and WRI1 detected in *pSPF1:SPF1-GFP* seeds. DAP, days after pollination. SPF1 and WRI1 proteins were detected using antibodies against GFP and WRI1, respectively. 12S globlin serving as the loading control. (J) Protein levels of SPF2 and WRI1 detected in *pSPF2:SPF2-GFP* seeds. DAP, days after pollination. SPF2 and WRI1 proteins were detected using antibodies against GFP and WRI1, respectively. 12S globlin serving as the loading control.

### SPF1 and SPF2 interact with WRI1

We previously showed that SIZ1-mediated SUMOylation regulates the stability of the WRI1 protein and hypothesized that WRI1 may be a substrate for the SUMO proteases SPF1 and SPF2 (Huang et al., 2025). SUMOylation is reversible, and SUMO proteases are known to function in deSUMOylation (Yates et al., 2016; Castro et al., 2018). To explore whether WRI1 is regulated by SUMO proteases, we investigated the potential interactions among WRI1, SPF1 and SPF2. We first employed the yeast two-hybrid system to verify their interactions and confirmed that WRI1 indeed interacts with SPF1 and SPF2 (Fig. 3A). To confirm the specificity of this interaction, we used the yeast two-hybrid system to screen for interactions between WRI1 and other SUMO proteases, such as ESD4, OTS1, OTS2, ULP1a, and ULP1b. The results showed that WRI1 does not interact with these SUMO proteases, indicating that the interaction between WRI1 and SPF1 and SPF2 is specific (Fig. 3A). To map the interaction domains, we divided the WRI1 protein into three segments based on its predicted domains: two AP2 domains and one TAD domain (transcription activation domain). Using SPF1 and SPF2 as bait and different WRI1 domains as prey (Fig. 3B), we found that only WRI1 truncations containing the highly conserved TAD domain interacted with SIZ1 (Fig. 3B). To confirm these interactions in vivo, we performed a bimolecular fluorescence complementation (BiFC) assay. YFP fluorescence was reconstituted when SPF1-YFP-C or SPF2-YFP-C was co-expressed with WRI1-YFP-N, and the SPF1-WRI1 and SPF2-WRI1 complexes were localized in the nucleus (Fig. 3C). Furthermore, we validated the in vivo association using a co-immunoprecipitation (co-IP) assay. MYC-SPF1 and MYC-SPF2 were co-immunoprecipitated by WRI1-GFP, supporting the specificity of the interaction between SPF1/2 and WRI1 (Fig. 3D).

**Figure 3.**
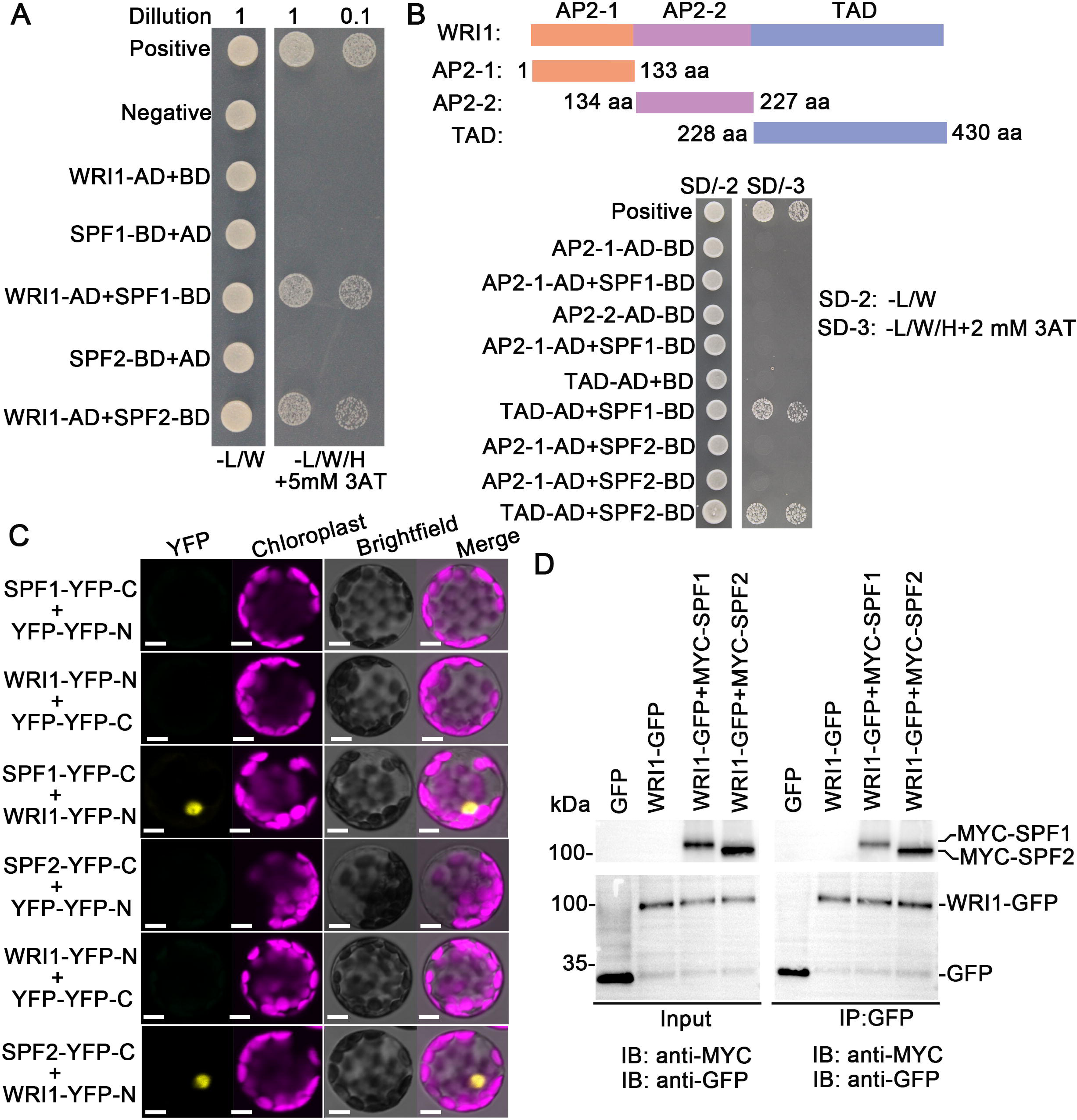
SPF1 and SPF2 interact with WRI1. (A) Confirmation of the interaction among SPF1, SPF2 and WRI1 through a yeast two-hybrid assay. Positive control: pGADT7-T + pGBKT7-53; negative control: pGADT7-T + pGBKT7-lam. WRI1-AD represents WRI1 fused to GAL4-AD; BD denotes GAL4-BD vectors without insertion; SPF1-BD and SPF2-BD signifies SPF1 and SPF2 fused to GAL4-BD; AD denotes GAL4-AD vectors without insertion; –L/W indicates selective medium Trp and Leu; – L/W/H denotes selective medium lacking Trp, Leu, and Ade. 3AT signifies 3-amino-1,2,4-triazole. (B) Identification of the domains in WRI1 required for their interaction using a yeast two-hybrid analysis. AP2-1/2-2: AP2 domains in WRI1. TAD: Region containing a transcriptional activation domain in WRI1. (C) Visualization of the interaction among SPF1, SPF2 and WRI1 in an *in vivo* Bimolecular Fluorescence Complementation (BiFC) assay in *Arabidopsis* protoplasts. Bars = 5 μm. (D) Validation of the interaction among SPF1, SPF2 and WRI1 interaction in an *in vivo* coimmunoprecipitation assay. Total protein extracts from transgenic plants carrying both 35S:MYC-SPF1 and UBQ10:WRI1-GFP or both 35S:MYC-SPF2 and UBQ10:WRI1-GFP were immunoprecipitated with immobilized anti-GFP antibody. The proteins from crude lysates (left) and immunoprecipitated proteins (right) were detected using anti-MYC or anti-GFP antibody.

### SPF1 and SPF2 regulates WRI1 protein stability by modulating its degradation

Our prior research has demonstrated that SUMOylation of WRI1 stabilizes the protein by modulating its degradation (Huang et al., 2025). Based on these findings, we hypothesize that SPF1 and SPF2, which influence the protein level of WRI1, may also exert their effects through a similar mechanism involving regulation of WRI1 degradation. To investigate this potential regulatory relationship, we firstly examined *WRI1* gene expression levels in 12 DAP seeds. The WRI1 gene expression level showed no difference across different SPF1/2 genotypes (data not shown), indicating that SPF1/2 may influence WRI1 translation or degradation. To explore how SPF1/2 affects WRI1 stability, 12 DAP seeds of wild type, *spf1-2spf2-1*, and *SPF1-OESPF2-OE* were subjected to treatment with the protein synthesis inhibitor CHX or the protease inhibitor MG132. Under control conditions, the protein level of WRI1 was significantly increased in the *spf1-2spf2-1* mutant, while it was significantly decreased in *SPF1-OESPF2-OE* seeds, indicating that SPF1/2 affects the stability of WRI1 protein (Fig 4A,4B). Interestingly, CHX treatment did not alter the overall trend in WRI1 protein levels across different SPF1/2 genotypes. (Fig 4C,4D). In contrast, MG132 treatment effectively restored WRI1 protein levels in *SPF1-OESPF2-OE* to those observed in wild-type plants (Fig 4E,4F). These results indicate that SPF1 and SPF2 regulates WRI1 protein stability by modulating its degradation.

**Figure 4.**
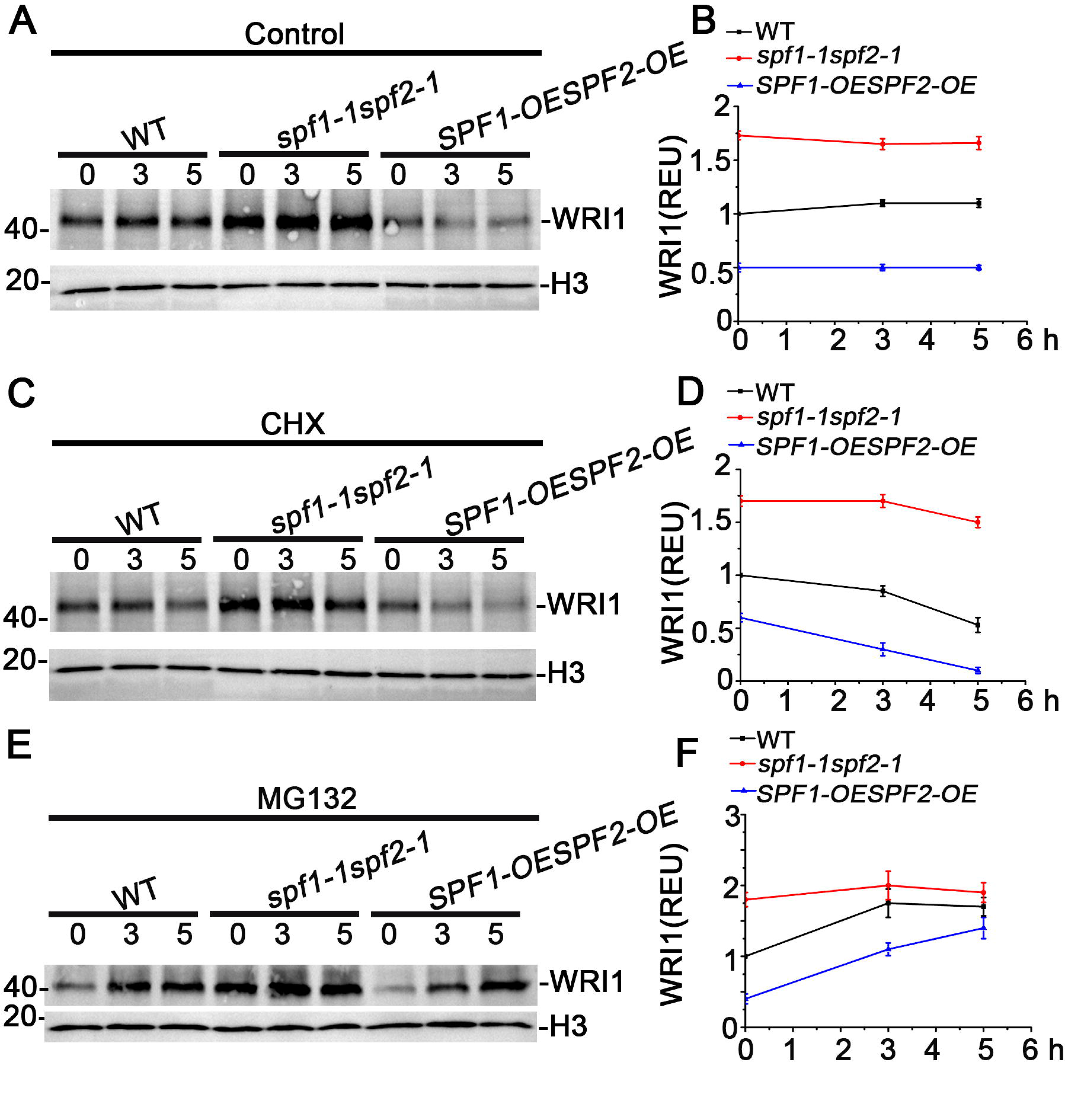
SPF1 and SPF2 regulates WRI1 protein stability by modulating its degradation. (A). Assessment of WRI1 protein levels in WT, *spf1-1spf2-1* and *SPF1-OESPF2-OE* plants under normal temperature conditions. (B) Quantified protein levels in (A) with three biological repeats are shown. (C) Measurement of WRI1 protein levels in WT, *spf1-1spf2-1* and *SPF1-OESPF2-OE* seeds. 12 DAP seeds were treatments with CHX for 3 or 5 hours. (D). Quantified protein levels in (C) with three biological repeats are shown. (E) Measurement of WRI1 protein levels in WT, *spf1-1spf2-1* and *SPF1-OESPF2-OE* seeds. 12 DAP seeds were treatments with MG132 for 3 or 5 hours. (F). Quantified protein levels in (E) with three biological repeats are shown. (B, D, F) The relative level of WRI1 accumulation as presented in relative expression units (REUs) is calculated based on the formula (WRI1^t^/H3^t^)/(WRI1°/H3°). in which “WRI1” and “H3” denote the digitized band intensities of WRI1 or H3 in the respective samples collected at time 0 or at the indicated time (T) after treatment. Error bars represent SD (n = 3).

### SPF1 and SPF2 affect the stability of the WRI1 protein via deSUMOylation

SUMO proteases play a crucial role in the dynamic regulation of SUMOylation (Yates et al., 2016). Therefore, we investigate whether the WRI1 protein undergoes deSUMOylation in an SPF1 and SPF2-dependent manner. In vivo SUMOylation analysis revealed that SPF1 and SPF2 can suppress the SUMOylation of WRI1 (Fig. 5A, 5B), with the combined presence of SPF1 and SPF2 almost abolishing the SUMOylation of WRI1 (Fig. 4C). This finding suggests that SPF1 and SPF2 may regulate WRI1 SUMOylation through their SUMO protease activity. To explore this hypothesis, we generated MYC-SPF1 and MYC-SPF2 plants with defective SUMO protease activity based on the analysis of protein sequence conservation (Kong et al., 2017). As expected, in vivo SUMOylation analysis showed that MYC-SPF1^C577S^ and MYC-SPF2^C485S^ had no effect on WRI1 SUMOylation (Fig. 5D, 5E). To corroborate that SPF1 and SPF2 regulate WRI1 SUMOylation, we also examined WRI1 SUMOylation in the *WRI1-GFP/spf1-2*, *WRI1-GEP/spf2-1*, and *WRI1-GEP/spf1-2spf2-1* plants. In vivo SUMOylation analysis revealed that WRI1 SUMOylation was enhanced in *WRI1-GFP/spf1-2* and *WRI1-GFP/spf2-1*, particularly in *WRI1-GEP/spf1-2spf2-1*(Fig. 5F-5H), suggesting that the WRI1 protein can undergo deSUMOylation in an SPF1 and SPF2-dependent manner.

**Figure 5.**
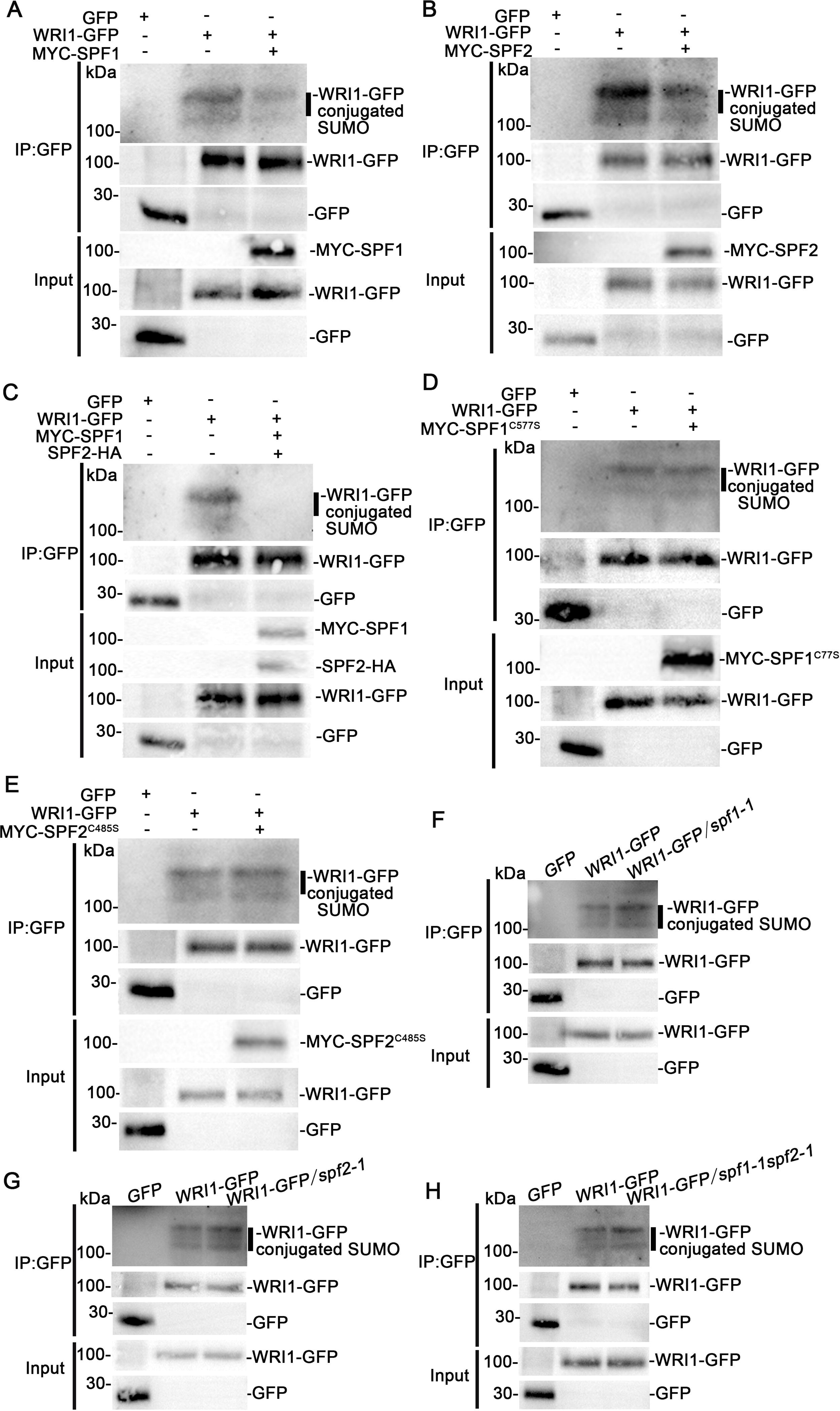
SPF1 and SPF2 affect the stability of the WRI1 protein via deSUMOylation. (A) Assessment of the effect of SPF1 or SPF2 on the SUMOylation modification of WRI1 in plants. The 12 DAP (day after pollination) seeds of SPF1-MYC/WRI1-GFP were used to analysis. (C). Assessment of the effect of SPF1 and SPF2 on the SUMOylation modification of WRI1 in plants. The 12 DAP (day after pollination) seeds of *SPF1-MYC/WRI1-GFP* were used to analysis. (D-E) Assessment of the effect of SPF1^C577C^ or SPF2^C485S^ on the SUMOylation modification of WRI1 in plants. The 12 DAP (day after pollination) seeds of SPF1-MYC/WRI1-GFP were used to analysis. (F) Assessment of WRI1 SUMOylation status in developing seeds of the WRI1-OE/spf1-1 at 12 DAP. (F) Assessment of WRI1 SUMOylation status in developing seeds of the *WRI1-OE/spf2-1* at 12 DAP. (F) Assessment of WRI1 SUMOylation status in developing seeds of the *WRI1-OE/spf1-1spf2-1* at 12 DAP.

### The deSUMOylation of WRI1 promotes its interaction with BPM2 and its ubiquitination

To assess how SUMOylation influences WRI1 stability, we analyzed transgenic lines expressing *pWRI1:WRI1-GFP* with comparable transcript levels in WT, *spf1-2spf2-1*, and *MYC-SPF1SPF2-HA* backgrounds. Analysis of GFP signals in embryonic cells demonstrated that WRI1-GFP accumulation was significantly reduced in *MYC-SPF1SPF2-HA* seeds compared to WT, whereas *spf1-2spf2-1* seeds exhibited markedly enhanced WRI1-GFP accumulation (Fig. 6A–6C). Since WRI1 is known to interact with BTB/POZ-MATH227 (BPM), an adaptor for CULLIN3-based ubiquitin E3 ligase that promotes ubiquitination and proteasomal degradation (Chen et al., 2013), we hypothesized that SUMOylation status modulates this regulatory pathway. Given the crosstalk between SUMOylation and ubiquitination (Rott et al., 2017), Specifically, we targeted BPM2 and performed co-immunocoprecipitation assays in *spf1-2spf2-1* and *MYC-SPF1SPF2-HA* plants to assess its interaction with WRI1. The analysis revealed that SPF1 and SPF2 distinctly enhance the interaction between WRI1 and BPM2, while a weaker interaction was observed in *spf1-2spf2-1* plants (Fig. 6D). Subsequently, we immunoprecipitated WRI1 proteins from 12 DAP seeds using an anti-GFP antibody and examined their ubiquitination states with an anti-ubiquitin antibody via immunoblotting. The western blot assay indicated that SPF1 and SPF2 substantially promote the ubiquitination of WRI1, whereas there is notably lower ubiquitinated WRI1 in *spf1-2spf2-1* compared to wild-type plants (Fig. 6E). These results collectively demonstrate that WRI1 deSUMOylation facilitates its interaction with BPM2 and subsequent ubiquitination, thereby influencing its stability.

**Figure 6.**
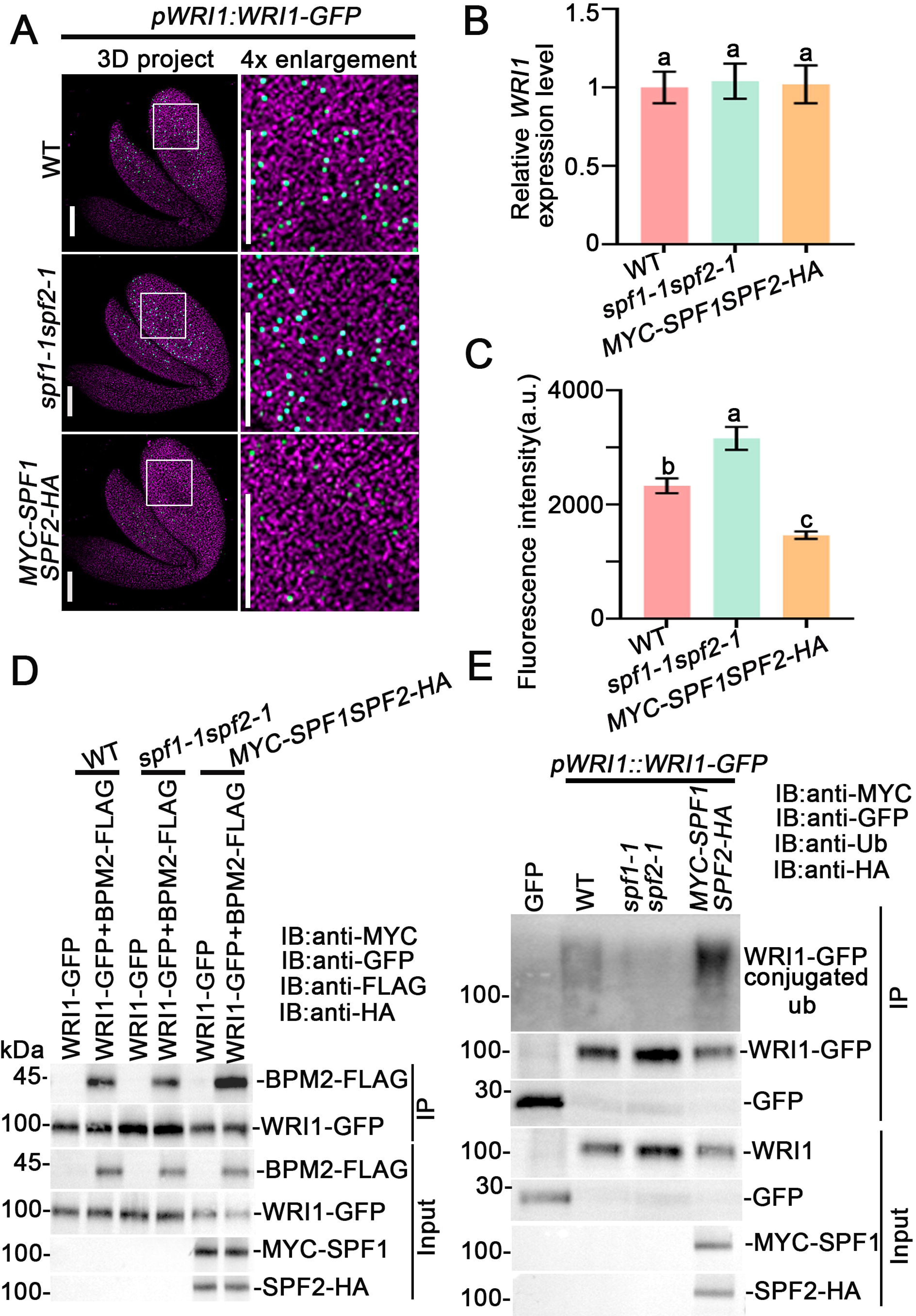
The deSUMOylation of WRI1 promotes its interaction with BPM2 and its ubiquitination. (A) Levels of WRI1-GFP protein accumulation in seeds. GFP fluorescence (green) was observed in 12 DAP *WRI1-GFP/*WT, *WRI1-GFP/spf1-1spf2-1* and *WRI1-GFP/SPF1-OESPF2-OE* embryos. Bars= 50 μm. (B) The WRI1 transcript levels in *WRI1-GFP/*WT, *WRI1-GFP/spf1-1spf2-1* and *WRI1-GFP/SPF1-OESPF2-OE* used in (A). *UBQ10* was used as a reference gene. (B) Measurement of the GFP fluorescence intensity in 12 DAP *WRI1-GFP/*WT, *WRI1-GFP/spf1-1spf2-1* and *WRI1-GFP/SPF1-OESPF2-OE*embryos. Chloroplast fluorescence was shown as magenta. Bars=50 μm. Ten embryos (n=10) of each treatment were used for GFP intensity calculating. The fluorescence signal were calculated in at least 60 nuclei from each embryo, including 20 nuclei each from the two cotyledons and the radicle. Fluorescence intensity labeled with the same letter are not significantly different, as determined by Tukey’s HSD test (P< 0.05). (D) Influence of deSUMOylation on the interaction between WRI1 and BPM2. The total protein was extracted from protoplasts transformed with WRI1-GFP or WRI1-GFP+BPM2-FLAG into WT, *spf1-1spf2-1*, and *SPF1-OESPF2-OE*. Immunoprecipitated with anti-GFP antibody was followed by detection of immunoprecipitated proteins using anti-GFP or anti-FLAG antibodies.

### SPF1 and SPF2 suppress fatty acid biosynthesis in seeds by deSUMOylation of WRI1

To further investigate whether SPFs and WRI1 function together in seed filing, we proposed the hypothesis that SPFs might influence seed filing by affecting WRI1 stability through deSUMOylation. To test this hypothesis, we assessed the genetic interaction between WRI1 and SPFs in regulating seed mass and seed fatty acid content. Specifically, we crossed the *spf1-2 spf2-1* double mutant with the *wri1-4* and analyzed the seed mass and fatty acid content in WT, *wri1-4*, *spf1-2 spf2-1*, and the *spf1-2spf2-1wri1-4* triple mutant plants. The results showed that the seed mass and fatty acid content of the *spf1-2spf2-1wri1-4* triple mutant were phenotypically similar to those of the *wri1-4* mutant seeds (Fig. 7A–7C), thereby confirming that SPF1 and SPF2 act in the WRI1 pathway during seed filing. Importantly, overexpression of WRI1 successfully rescued the phenotypes of *SPF1-OESPF2-OE* seeds (Fig. 7A–7C), whereas *SPF1-OE SPF2-OE* did not restore the seed mass or fatty acid content of the *wri1-4* mutant (Fig. 7A–7C). These findings strongly suggest that SPF1 and SPF2 regulate seed size and fatty acid content in a WRI1-dependent manner. Furthermore, the seed phenotypes of *WRI1-OE/spf1-2spf2-1* plants were comparable to those of *WRI1-OE* seeds (Fig. 7A–7C), thus highlighting that SPF1 and SPF2 suppress fatty acid biosynthesis in seeds through WRI1 deSUMOylation.

**Figure 7.**
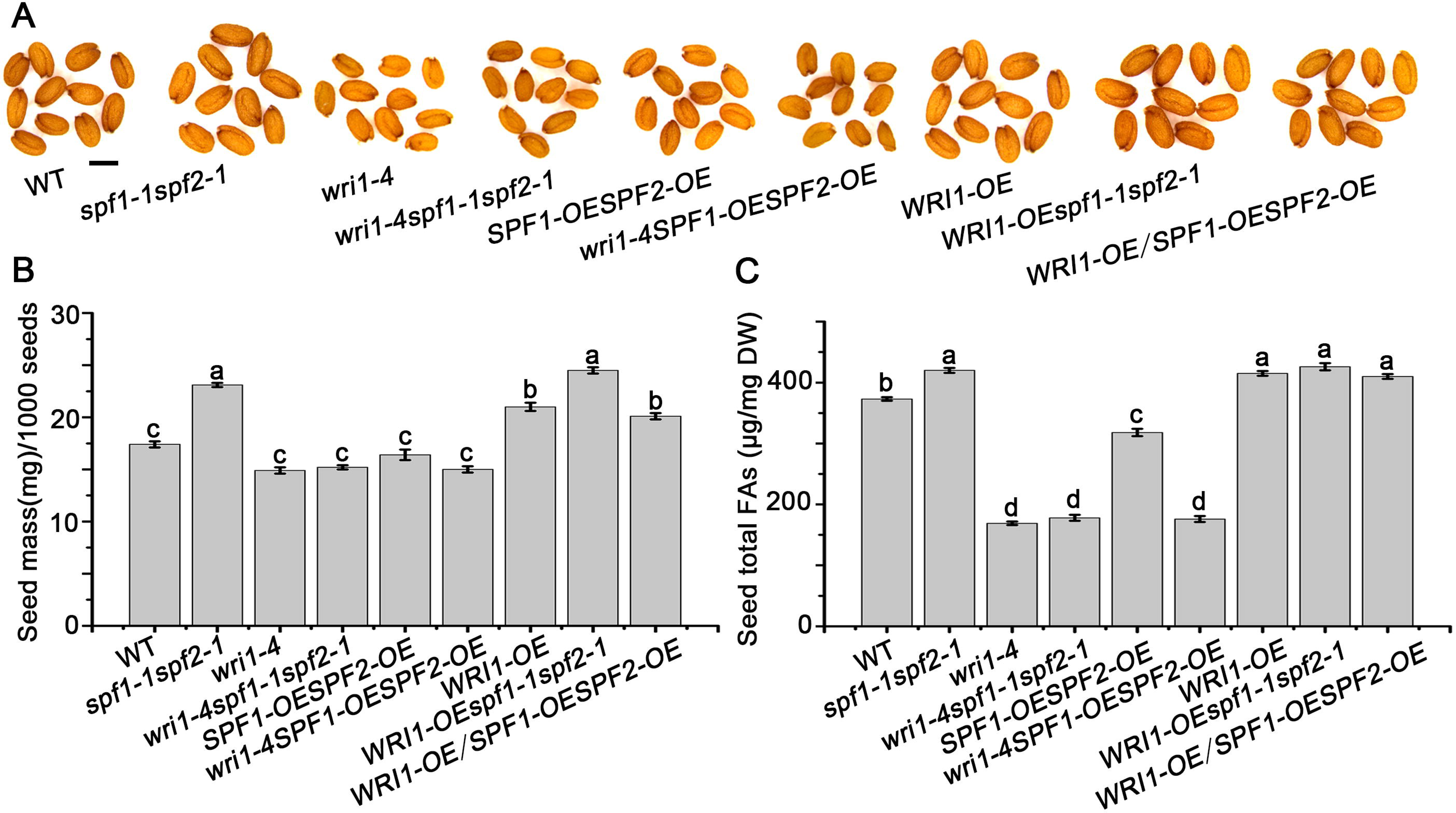
SPF1 and SPF2 suppress fatty acid biosynthesis in seeds by deSUMOylation of WRI1. (A) Stereoscopic examination of mature seed phenotypes in WT, *wri1-4*, *WRI1-OE*, *spf1-1*, *spf2-1*, *spf1*-*1spf2*-*1wri1*-4, *WRI1*-OE/*spf1*-*1spf2*-1, *SPF1*-*OESPF2*-*OE*, *SPF1*-*OESPF2*-*OE/wri1-4,* and *WRI1-OESPF1-OESPF2-OE* plants. Bars = 500 μm. (B) Measurement of seed mass in WT, *wri1-4*, *WRI1-OE*, *spf1-1*, *spf2-1*, *spf1*-*1spf2*-*1wri1*-4, *WRI1*-OE/*spf1*-*1spf2*-1, *SPF1*-*OESPF2*-*OE*, *SPF1*-*OESPF2*-*OE/wri1-4,* and *WRI1-OESPF1-OESPF2-OE* plants. (C) Assessment of fatty acid contents in seeds from WT, *wri1-4*, *WRI1-OE*, *spf1-1*, *spf2-1*, *spf1*-*1spf2*-*1wri1*-4, *WRI1*-OE/*spf1*-*1spf2*-1, *SPF1*-*OESPF2*-*OE*, *SPF1*-*OESPF2*-*OE/wri1-4,* and *WRI1-OESPF1-OESPF2-OE* plants. The data shown represent means ± standard deviation (SD) (N = 3), with each assay involving 1000 seeds per biological replicate. Seed mass values labeled with the same letter are not significantly different, as determined by Tukey’s HSD test (P< 0.05).

## Discussion

The dynamic regulation of WRI1 protein stability is central to lipid biosynthesis during seed development. The regulation of seed filing and fatty acid biosynthesis is a complex process involving intricate transcriptional and post-translational mechanisms. In this study, we uncovered a novel regulatory pathway in which SPF1/SPF2-mediated deSUMOylation promotes WRI1 degradation via the ubiquitin-proteasome system, while WRI1 reciprocally suppresses *SPF1/SPF2* transcription to stabilize its own protein levels. Our findings not only shed light on the dynamic interplay between SUMOylation and protein stability but also highlight the critical role of feedback regulation in maintaining WRI1 homeostasis.

SPF1 and SPF2 are pivotal SUMO proteases in plant development, particularly in *Arabidopsis* fertility (Kong et al., 2017; Liu et al., 2017; Verma et al., 2021). However, the regulatory mechanism of *SPF1* and *SPF2* in development has been scarcely studied. In this investigation, we discovered that SPF1 and SPF2, previously identified as essential for fertility in *Arabidopsis* (Liu et al., 2017), also suppress seed filling. Seed filling is an important stage of seed development during which embryos gradually accumulate seed storage proteins and oils, both of which account for 30%– 40% of dry seed matter (Baud et al., 2008). Seed oil is mainly accumulated from 6 DAP and stored in embryos in Arabidopsis (Baud et al., 2008). The genetic results showed that SPF1 and SPF2 suppress seed oil accumulation (Fig.1A-1C), which is highly consistent with its strong expression in developing embryos from 6 and 16 DAP (Fig. 2A-2D).. Embryo cells undergo cell division and expansion, thereby affecting the sizes of seeds and embryos, the weight of seeds, and the accumulation of seed storage reserves (Baud et al., 2008). Our results showed that the SPF1 and SPF2 mutation resulted in an increase in the seed weight and oil content, while *SPF1* and *SPF2* over expression exhibit a reduction in the seed weight and oil content, which corresponds to the seeds phenotype. Moreover, In the *spf1*, *spf2 and spf1-2spf2-1* mutants, the filling process is delayed accompanied by a significant decrease in seed oil content (Fig. 1D), indicated that SPF1 and SPF2 may participated in regulating seed filling..The LAFL genes are mater transcription factors in seed filling, involved in the accumulation of storage materials such as lipids and storage proteins in seeds (Boulard et al., 2017). Although we cannot definitively conclude whether the LAFL genes are direct targets of SPF1 and SPF2, this result also suggests the requirement of SPF1 and SPF2 for seed filling. Thus, SPF1 and SPF2 might regulate seed storage protein accumulation, with some regulatory factors potentially playing a pivotal role in this process as substrates of SUMO proteases SPF1 and SPF2.

The AP2 transcription factor WRI1 plays a critical role as a key regulator of fatty acid synthesis in Arabidopsis (Cernac & Benning, 2004; Maeo et al., 2009; Ma et al., 2013). SIZ1-mediated SUMOylation modification maintains WRI1 protein stability by regulating ubiquitination (Huang et al., 2025). SUMOylation modification is reversible post-translational modifications of proteins. We found that SPF1 and SPF2 physically interacts with WRI1 (Fig. 3A, 3B). Subsequently, we used the yeast two-hybrid system to verify the interactions between WRI1 and other known SUMO proteases such as ESD4, OTS1, OTS2, ULP1a, and ULP1b in Arabidopsis thaliana(data not shown). The results showed that WRI1 did not interact with these SUMO proteases,indicating that the interaction between SPF1/2 and WRI1 is specific. However, we cannot currently rule out the possibility that WRI1 interact with these SUMO proteases in vivo.

In *Arabidopsis*, the SUMO proteasome has been reported to influence substrate abundance through its protease activity. For instance, OTS1/OTS2 deSUMOylatee JAZ and DELLA proteins, modulating jasmonic acid and gibberellin signaling, respectively, while SPF1 and SPF2 deSUMOlytion FLC and EDA9, interfering with flowering time and fertility, respectively (Kong et al., 2017; Liu et al., 2017; Srivastava et al., 2018; Conti et al., 2008; Conti et al., 2014). Overexpression of SPF1 and SPF2 reduces WRI1 protein levels, while mutations in *SPF1* and *SPF2* increase WRI1 protein levels (Fig. 4A, Fig. 6A). Notably, the gene expression levels of WRI1 show no significant differences across different genotypes. These results suggest that SPF1 and SPF2 SUMO proteases likely influence WRI1 protein stability rather than its transcriptional regulation. Our study further clarifies how SPF1 and SPF2 regulates WRI1 stability through protein synthesis and degradation assays. After treatment with the protein synthesis inhibitor cycloheximide (CHX) in 12 DAP seeds of the *spf1-1/spf2-1* and *SPF1-OESPF2-OE* overexpressing seeds. The overall variation trend in WRI1 protein levels in these plant seeds was similar to that under control conditions, indicating that SPF1 and SPF2 do not affect the synthesis of the WRI1 protein. However, after treatment with the proteasome inhibitor in seeds of these plants, the WRI1 protein level in SPF1-OE SPF2-OE overexpressing seeds was restored to that observed in wild-type (WT) seeds (Fig. 4). This result suggests that SPF1/2 influences the stability of the WRI1 protein by affecting its proteasomal degradation rather than its synthesis. These findings align with the established role of SUMOylation in stabilizing WRI1 protein levels (Huang et al., 2025).

SPF1 and SPF2 exhibit deSUMOylation activity toward WRI1. When both SUMO proteases are present, the SUMOylation of WRI1 is nearly abolished, whereas depletion of either SPF1 or SPF2 markedly enhances WRI1 SUMOylation (Fig. 5). These findings align with seed phenotypes (Fig. 1), suggesting that SPF1 and SPF2 are not functionally redundant but instead act synergistically to regulate oil accumulation during seed filling. Moreover, the mutant forms of SPF1 and SPF2 proteins with SUMO protease activity site mutations are unable to deSUMOylate the WRI1 protein and do not affect the stability of the WRI1 protein (Fig. 5E-5F). These observations further demonstrate that SPF1 and SPF2 mediate the deSUMOylation of the WRI1 protein and thereby influence its stability.

Previous studies have revealed that WRI1 can undergo degradation mediated by the CUL3-mediated ubiquitin proteasome pathway involving the BPMs protein (Chen et al., 2013). Our results demonstrate that the presence of the SUMO proteases SPF1/SPF2 promotes the interaction between WRI1 and BPM2 (Fig. 6D). Additionally, we observed that the ubiquitination levels of WRI1 were significantly higher in *SPF1-OESPF2-OE* 12 DAP seeds, whereas they were notably lower in *spf1-2spf2-1* mutants compared to the wild type (Fig. 6E). These findings strongly suggest that the deSUMOylation of WRI1 promotes its interaction with BPM proteins, subsequently enhancing its ubiquitination. Furthermore, genetic relationship analysis revealed that WRI1 acts downstream of SPF1 and SPF2 to regulate seed filling and fatty acid biosynthesis (Fig.7A-7C). We also observed that *SPF1-OESPF2-OE* was unable to rescue the seed phenotype of *wri1-4* mutants, including seed mass and fatty acid content, whereas overexpression of WRI1 successfully rescued the seed phenotype of *SPF1-OESPF2-OE* (Fig. 7A-7C). Therefore, SPF1 and SPF2 regulate WRI1 protein stability by mediating its deSUMOylation, thereby influencing seed filling.

In summary, this study clarifies the molecular mechanism by which deSUMOylation and ubiquitination crosstalk regulate seed filling. SPF1 and SPF2 mediate the deSUMOylation of WRI1, thereby regulating the ubiquitination of the transcription factor WRI1 to regulate the stability of WRI1.

## Materials and Methods

### Plant Materials and Growth Condition

The *Arabidopsis* T-DNA insert mutant line *spf1-1* (SALK_040576C, *spf2-1* (SALK_023493C) and *wri1–4* (SALK_ 008559) were obtained from the Arabidopsis Biological Resource Center in the USA (http://signal.salk.edu) All genotypes used in this study are of the Col-0 background. To generate the transgenic plants expressing *pSPF1:GUS* and *pSPF2:GUS*, 2.5 kb promoter of *SPF1* and *SPF2* was polymerase chain reaction (PCR)-amplified and inserted into the binary vector *pCAMBIA-1300221* respectively through the restriction endonuclease sites Hind III and BamH I (Huang et al., 2021). The *SPF1-GFP (SPF1-OE)* and *SPF2-GFP (SPF2-OE)* transgenic lines were generated as follows. The *SPF1* or *SPF2* coding region was amplified and inserted into the vector *pCAMBIA-1300-GFP* (driven by the *UBQ10* promoter; Huang et al., 2021). To generate the *pSPF1:SPF1-GFP* and *pSPF2:SPF2-GFP* plant, the full-length coding sequence of SPF1 and SPF2 was PCR-amplified and cloned into *pCAMBIA1300* driven by the 2.5kb native promoter respectively. To generate the *MYC-SPF1* and *MYC-SPF2* plant, the full-length coding sequence of SPF1 and SPF2 was PCR-amplified and cloned intothe vector *pCANG* driven by the *Cauliflower Mosaic Virus 35S Promoter* (CaMV 35S). To generate the *SPF2-HA* plant, the full-length coding sequence of SPF2 was PCR-amplified and cloned intothe vector *pBA002* driven by the CaMV 35S promoter. To generate the *BPM2-FLAG* plant, the full-length coding sequence of *BPM2* was PCR-amplified and cloned intothe vector *pBA002* driven by the CaMV 35S promoter. The resulting constructs in binary vectors were transformed into Agrobacterium and introduced into WT by floral dipping. Seeds were surface-sterilised and stratified at 4°C for 48 h in darkness before sowing on standard Murashige and Skoog (MS) plates supplemented with 1.5% sucrose and 0.7% agar. Arabidopsis plants were grown at 22°C under long-day conditions.

### Histochemical staining

Histochemical staining to assess GUS activity in homozygous transgenic plants and developing embryo was conducted following the method originally described by Jefferson et al. (1987) with some modification. A GUS stock solution, comprising 0.05 M NaPO_4_ buffer (pH 7.0), 5mM K_3_Fe(CN)_6_, 5mM K_4_Fe(CN)_6_, and 10 mM X-glucuronide, was prepared as previously outlined (Stahl et al., 2009). Developing seeds at various stages were dissected from siliques, immersed in the GUS staining solution, and incubated at 37°C in the dark for 2 to 8 hours, depending on the experimental requirements. After incubation, the seeds were rinsed with 75% ethanol and precleared in Hoyers’ solution for 1 to 12 hours, depending on their developmental stage and experimental needs. The stained embryos were then examined under a differential interference contrast stereoscope (Leica), as described by Huang et al. (2022). Each experiment included three biological replicates, with at least 20 embryos of each genotype stained and examined.

### Confocal Microscopy

For examination of homozygous transgenic plants through confocal microscopy, developing seeds at various stages were dissected from siliques, rinsed in distilled water, and mounted in water, following the protocol outlined by Huang et al. (2022) with some modification. By using the Zeiss LSM 800 laser scanning microscope, the subcellular distribution of GFP fusion protein can be observed by exciting at 514 nm and detecting emission between 530-600 nm, depending on whether 3D stacking technology is used as needed. Chlorophyll autofluorescence was concurrently captured by exciting at 488 nm and detecting emission between 650-700 nm, and images from the GFP, chlorophyll, and bright field channels were merged for analysis.

### Fluorescence quantification

The quantification of fluorescence intensity for WRI1-GFP was conducted the methodology described by Wilma van Esse et al. (2011) using FIJI software (Schindelin et al., 2012). The area in pixels and raw integrated density of the fluorescence signal were measured in at least 60 nuclei from each embryo, including 20 nuclei each from the two cotyledons and the radicle. Fluorescence intensity (arbitrary units) was calculated by dividing the raw integrated density by the area in pixels of the fluorescence signal.

### Bifluorescence complementation (BiFC) assays

Rosette leaves from 4-week-old *Arabidopsis* plants were used for protoplast transformation, employing the protocol established by Jen Sheen’s laboratory, as previously outlined (Yoo et al., 2007). For the BiFC experiment, the WRI1-YFP-N vector was constructed by integrating the WRI1 coding region into the pSPYNE plasmid, while the SPF1-YFP-C and SPF2-YFP-C vector was generated by inserting the SPF1 and SPF2 coding region into the pSPYCE plasmid (Walter et al., 2004). The transformed protoplasts were cultured for 48 hours at 23°C in the dark. The subcellular localization of YFP fusion proteins was examined using a Zeiss LSM 710 laser scanning microscope with excitation at 514 nm and emission detection between 530–600 nm. Chlorophyll autofluorescence was concurrently captured by exciting at 488 nm and detecting emission between 650-700 nm, and the images from the YFP, chlorophyll, and bright field channels were merged for analysis.

### Yeast Two-Hybrid Assays

Yeast two-hybrid assays were performed using the Matchmaker GAL4-based Two-Hybrid System 3 (Clontech), following the manufacturer’s guidelines. Full-length SPF1, SPF2, ESD4, OTS1, OTS2, WRI1, ULP1a, ULP1b and the three segmented structural domains of WRI1 were integrated into the Gal4 DNA binding domain (BD) and the Gal4 activation domain (AD). These plasmids were co-transformed into the yeast strain Y2HGold (Clontech). Equal quantities of yeast cells from each transformed strain were then plated on synthetic defined (SD)-Leu/-Trp and SD-Leu/-Trp/-His/ selective medium and incubated at 30 °C for 2–4 days.

### RNA extraction and gene expression analysis

Total RNA was extracted from seeds at various developmental stages using a Plant RNA Kit (PROMEGA, Madison, WI, USA) following the manufacturer’s instructions. The RNA was used to synthesize cDNA with oligo (dT) primers via reverse transcription (RT). Specifically, 1 g of total RNA with oligo (dT) primer was heated at 70 °C for 10 min, then cooled, and combined with MMLV-RT SPCL reverse transcriptase (Invitrogen, Waltham, MA, USA) for 1 h at 42 °C. The *UBQ10* (At4g05320) served as the reference gene (Czechowski et al., 2005). The qPCR was conducted using a C1000 Thermal Cycler (CFX96 Real-Time System, Bio-Rad, Hercules, CA, USA). Expression levels were determined and analyzed using the 2^−ΔΔCq^ method with Bio-Rad CFX Manager software (version 2.1). Each experiment included three technical replicates of the same sample and three biological replicates from distinct samples collected at different times.

### Immunoprecipitation

Proteins were extracted from the 12 DAP siliques of related transgenic plants in extraction buffer (50 mM Tris–HCl, pH 7.4, 150 mM NaCl, 2 mM MgCl2, 20%glycerol, and 0.1% Nonidet P-40) containing protease inhibitor cocktail (Roche, Basel, Switzerland). After centrifugation at 13,000 g for 12 min, the supernatant was incubated with GFP-Trap at 4 ℃ for 2 h. The beads were centrifuged and washed 4 times with washing buffer (50 mM Tris–HCl, pH7.4, 150 mM NaCl, 2 mM MgCl2, 10% glycerol, and 0.01% NP-40). Proteins were eluted with SDS sample buffer and detected with anti-MYC (TransGen, Beijing, China; HT101), anti-HA (TransGen, Beijing, China; HT301-01), anti-FLAG (TransGen, Beijing, China; HT201-01), anti-GST (TransGen, Beijing, China; HT601-01), anti-SUMO1 (Abcam, Ab5316) antibodie and anti-Ubquitin (Proteintech, Cat No. 10201-2-AP).

### Total fatty acid and seed storage protein quantification

Total fatty acids from corresponding dry seeds were extracted and quantified as previously described (Huang et al., 2025). Three biological replicates were performed in each experiment.

For seed storage protein quantification(SSP), seed storage protein was extracted and quantified as previously described with some modification (Hu et al., 2022). Dry seeds were collected and dried in 37℃ for 12 d to eliminate the impact of seed water content. Seeds were precisely weighed for 20.00 mg and finely ground in an electronic homogenizer. Next one milliliter SSP extraction buffer consist of 50 mM HEPES pH=7.5, 5 mM MgCl2, 5mM dithiothreitol (DTT), 1mM phenylmethylsulfonyl fluoride (PMSF), 1 mM ethylene diamine tetraacetic acid (EDTA) and 10% (v/v) Ethylene glycol was added to the ground seed powder. After centrifugation at 14,000 rpm for 10 min under 4℃, the supernatant of each sample was transferred to a new tube and then used for storage protein measurements. The remaining insoluble proteins were recovered by 1 M sodium hydroxide and measured as described above. In both assays, bovine serum albumin (BSA) was used for calibration.

### Immunoblotting and protein stability assays

To assess WRI1, SPF1 and SPF2 protein levels during seed filling, proteins were extracted from 4-, 6-, 8-, 10- 12-, 14- or 16-DAP seeds of *pSPF1:SPF1-GFP* and *pSPF2:SPF2-GFP* plants. The extraction buffer containing 50 mM Tris–HCl, pH 7.4, 150 mM NaCl, 2 mM MgCl_2_, 20% glycerol, and 0.1% NP-40, and a protease inhibitor cocktail (Roche, Basel, Switzerland) (Huang et al., 2022). Extracted proteins were separated by SDS–PAGE and analyzed via immunoblotting (IB) using anti-GFP (TransGen, Beijing, China; HT801-01), anti-WRI1 (designed by GeneScript) or anti-H3 (Abcam; ab176842) antibodies. For WRI1 stability assays, seeds were incubated in liquid MS medium with or without the addition of CHX (Sigma-Aldrich; 50 µM) or MG132 (Sigma-Aldrich; 50 µM) at 22 °C or 35 °C for 0, 3, or 5 hours, using DMSO was a control. After incubation, proteins were extracted with the same extraction buffer, separated by SDS–PAGE and analyzed via IB using anti-WRI1 or anti-H3 antibodies. The WRI1 antibody detect endogenous WRI1 protein at about 48.4 kD. Relative protein levels were quantified using ImageJ software by calculating the ratio of target protein to H3, standardized against control samples. Each experiment included at least three biological replicates.

## Statistical analyses

Statistically significant differences compared to the control (WT) were assessed using Student’s *t*-test. To evaluate gene expression and seed phenotype variations across different genotypes for a specific variable, we conducted one-way ANOVA followed by Tukey’s Honestly Significant Difference test (P <0.05). Groups denoted with the same letter indicate no significant differences. All statistical analyses were performed using SPSS 22.0 software (IBM Corp., Armonk, NY, USA).

## Accession numbers

Sequence data from this article can be found in the Arabidopsis information resource under the following accession numbers: *SPF1* (AT1G09730), *SPF2* (AT4G33620), *WRI1* (AT3G54320), *BPM2* (At3g06190).

## Acknowledgements

We thank the ABRC for kindly providing seeds used in this study. We thank Chao Yang from the South China Botanical Garden, Chinese Academy of Sciences, for providing the *SPF1* and *SPF2* overexpression plants and mutant plants.

## Author Contributions

S. Z. conceived this project and designed all research. R. H. performed most of the research. M. W., Y. C., B. F., M. Y., H. H., J, M., Q, L., W, L., Z, M., and L, Z. helped perform some experiments. Y. C. helped measure the fatty acid contents. S. Z., R. H., C. Y. and H, L. analyzed the data. S. Z. and R. H. wrote the article. S. Z. and R. H. revised the article.

